# Energy Landscape of the SARS-CoV-2 Reveals Extensive Conformational Heterogeneity

**DOI:** 10.1101/2021.05.11.443708

**Authors:** Ghoncheh Mashayekhi, John Vant, Abhishek Singharoy, Abbas Ourmazd

**Affiliations:** Department of Physics, University of Wisconsin Milwaukee, 3135 N. Maryland Ave, Milwaukee, WI, 53211, USA; School of Molecular Sciences, Center for Applied Structural Discovery, Arizona State University, Tempe, AZ, 85287, USA

## Abstract

Cryo-electron microscopy (cryo-EM) has produced a number of structural models of the SARS-CoV-2 spike, already prompting biomedical outcomes. However, these reported models and their associated electrostatic potential maps represent an unknown admixture of conformations stemming from the underlying energy landscape of the spike protein. As for any protein, some of the spike’s conformational motions are expected to be biophysically relevant, but cannot be interpreted only by static models. Using experimental cryo-EM images, we present the energy landscape of the spike protein conformations, and identify molecular rearrangements along the most-likely conformational path in the vicinity of the open (so called 1RBD-up) state. The resulting global and local atomic refinements reveal larger movements than those expected by comparing the reported 1RBD-up and 1RBD-down cryo-EM models. Here we report greater degrees of “openness” in global conformations of the 1RBD-up state, not revealed in the single-model interpretations of the density maps, together with conformations that overlap with the reported models. We discover how the glycan shield contributes to the stability of these conformations along the minimum free-energy pathway. A local analysis of seven key binding pockets reveals that six out them, including those for engaging ACE2, therapeutic mini-proteins, linoleic acid, two different kinds of antibodies, and protein-glycan interaction sites, switch conformations between their known apo- and holo-conformations, even when the global spike conformation is 1RBD-up. This is reminiscent of a conformational pre-equilibrium. We found only one binding pocket, namely antibody AB-C135 to remain closed along the entire minimum free energy path, suggesting an induced fit mechanism for this enzyme.

## Introduction

Intensive research, primarily by cryo-Electron Microscopy (cryo-EM) techniques, has established that the spike protein plays a critical role in the process of infection by the SARS-COVID family (*1,2*). The average ‘apo’ (ligand-free) and ‘holo’ (ligand-bound) conformations assumed by the spike protein are now known to near-atomic resolution (*1,3,4*). It is recognized that the spike protein exhibits structural variability (*1, 5*). However, access to an experimentally determined conformational path between the apo- and holo-end states would substantially elucidate the thermally accessible functionally relevant conformational motions of the spike protein (*6,7*).

As different molecular conformations populate different energy states, their conformational spectrum defines an energy landscape. This landscape is, in principle, multi-dimensional. Nonetheless, a two-dimensional (2D) representation described by the leading two conformational coordinates is commonly used to project the multidimensional free energy profiles on to a manifold of reduced dimensionality (*6–8*). Near equilibrium, biomolecular function corresponds to traversing the minimum-energy pathway connecting the initial and final states (*9–13*). For example, the path underlying cell recognition of the spike protein commences along the energy landscape of the apo state transitions to the holo state after binding the cell surface receptors, ending at the most probable conformations on the holo landscape. Here, we focus on the apo segment of the spike conformational pathway and draw inferences on its plausible binding mechanisms not amenable to a single-structure representation, or even microsecond-long molecular dynamics (MD) simulations of apo-state models.

The Receptor Binding Domain (RBD) of the apo spike protein exists primarily in two conformations. These “down” and “up” states occur with nearly equal probability (*1, 3*). In the down state, the ACE2 binding pocket is obscured, rendering membrane fusion essentially unfeasible. The nature of the pathway between the down and the up states of the RBD, and the transition rate between a hidden and an accessible ACE2 binding pocket are of central importance for understanding SARS-CoV2’s ability to hide vulnerable epitopes from the host’s immune system (*2*). Several studies have employed Molecular Dynamics (MD) simulations to investigate the transitions between these states, using static cryo-EM structures to characterize each endpoint (*14–19*). However, a single cryo-EM map often represents an ensemble of thermally accessible conformations (*7, 20, 21*). Brute-force MD simulations are thus expected to have an inherent model bias towards the selected starting structures. Enhanced sampling simulations have been successful in overcoming such bias (*22*), but are cumbersome.

Here, by combining all-atom MD simulations and cryo-EM data analysis, we determine the collection of conformations pertaining to the up state of the RBD. Our results show that in the apo-state, the spike protein assumes a broadly heterogeneous ensemble of nearly isoenergetic conformations. The heterogeneity of these structures can substantially exceed the differences between the average down and up structures inferred by standard cryo-EM analyses (*1, 3*). The local movements have a dramatic effect on key binding sites, offering fresh insights into how molecular recognition occurs at the spike’s ACE2, linoleic acid, antibodies-binding domains. Our approach offers a useful guide for future work on characterizing the most probable down to up transitions of the RBD, by integrating experimental cryo-EM data and MD simulations.

## Results

As described below, we extract the conformational energy landscape of the spike protein from experimental cryo-EM snapshots, and reconstruct 3D density maps along the minimum-free-energy pathway (MFEP) on this landscape. Molecular dynamics simulations are then employed to interpret the density information in all-atoms detail. Finally, we present the global and local conformations of the up-state of the apo spike protein, focusing on the conformational heterogeneity of key binding pockets.

### Conformational coordinates, energy landscape, and functional trajectory

We use geometric machine-learning (ManifoldEM) (*23*) to extract the manifold of conformational motions from cryo-EM images. This manifold is spanned by a set of orthogonal conformational coordinates (CC) (*6*). To determine the conformational changes along each coordinate, we compile a 3D movie of the density maps along each of those coordinates. (Details of the density movie generation scheme is provided in Methods). Fifty density maps were extracted along each of the two conformational coordinates. The nominal resolution of these maps varies from 3.2 to 4.4 Å. Molecular dynamics flexible fitting or MDFF was employed to construct molecular models from each of the maps (*24,25*), translating the density movies along CC1 and CC2 to molecular movies (Methods and Supplementary Movies 1 and 2).

By construction, the motions observed along CC1 play out in the XY plane, and motions observed for CC2 evolve along the Z-axis. Movement along CC1 corresponds to a global change in the RBD’s center of mass, which deforms orthogonally to the spike protein’s principal axis. Similarly, motion along CC2 captures a “projectile-like” motion of the RBD parallel to the spike protein’s principal axis. The functionally relevant conformational movements, however, involve a combination of CC1 and CC2 along the minimum-energy path on the conformational landscape (*9*). As detailed in Methods, we infer free-energy changes from the population of points on the CC1-CC2 conformational plane via the Boltzmann factor (fig. 1A) (*6*). The minimum free energy path on this CC1-CC2 landscape consists of a ‘horse-shoe’ shaped, essentially iso-energetic tube. Compiling the conformational movies along the MFEP reveals the functional motions of the spike protein (Supplementary Movie 3), because, near equilibrium, other conformational states are not significantly occupied.

**Figure 1:**
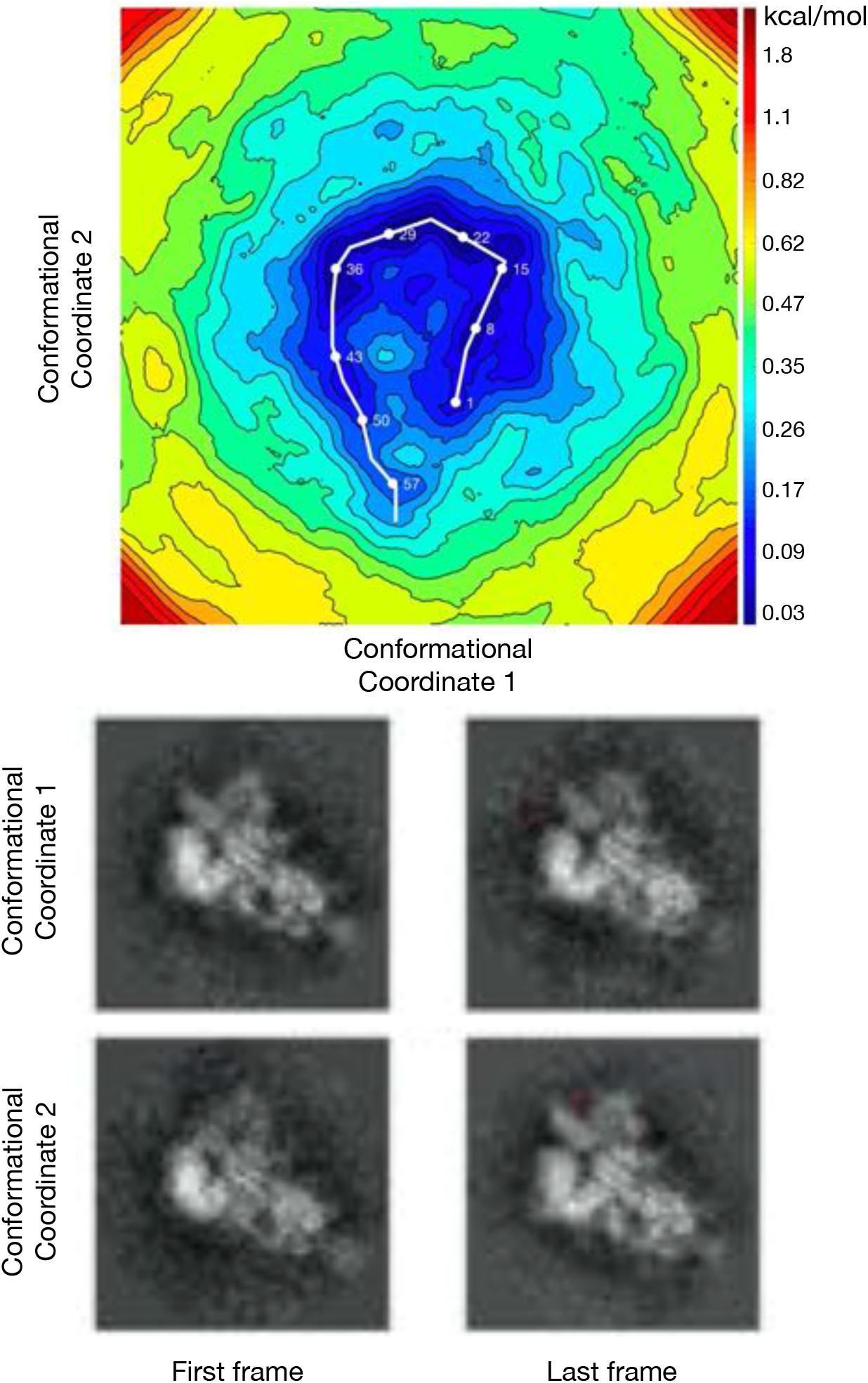
The energy landscape of the SARS-CoV-2 Spike Protein spanned by two conformational coordinates. Each conformational coordinate consists of a space of 50 binned electrostatic potential maps defining conformational changes in the space of these maps. The MFEP locations are traced with a white line representing the 59 MFEP locations found in this study. The numbering scheme used here is consistent with our analysis.

Previously reported 6VSB (*3*) and 6VYB (*1*) densities resemble ManifoldEM models pertaining to location no. 29 (CC1 21; CC2 31) and its close vicinity on the minimum free energy pathway in fig. 1A. (See Methods for relating density maps to points on the energy landscape.) Most of the states along the minimum free energy pathway are nearly isoenergetic, and thus populated with comparable probabilities. The pursuit of high-resolution structures, however, may preferentially select a subset of the high probability conformations present (*7*). key spike functions (e.g., antibody and receptor binding or epitope signaling) stem from the entire population of structures along the MFEP. The conformational heterogeneity associated with the states in this horse-shoe shaped reaction tube highlights the range of global and local conformations often subsumed by single-model representations, which ignore the underlying conformational energy landscapes. We already observe at the level of density changes, that the conformational spectrum along the MFEP of the apo up spike conformation can result in more open (CC1 25 → 31; CC2 28 → 32) and less open (CC1 17 → 20 and CC2 8 → 12) apo conformations. These conformations are as probable as those reported in the PDB (*1,3*). The variation in conformations of the spike protein is only partially apparent in a comparison between the reported static apo (*1, 3*), holo (*26–28*), and long-time MD simulation models of the spike (*29*). In contrast, we circumvent the uncertainties in inferring function from stationary snapshots (*6*), by compiling density movies along the MFEP to reveal the functional trajectory of the apo spike conformation.

### Global conformational changes along MFEP

A total of 59 points on MFEP were refined using a multi-grid MDFF procedure (*30,31*) to gauge the most probable conformational changes of the apo spike. Shown in Supplementary Movie 3, the set of conformations along the MFEP reveals deviations from the static apo structures at both ends of the horseshoe-shaped tube. As expected, models stemming from the MFEP resemble previously published open state structures 6VSB and 6VYB (*1,3*), with RMSD ranging from 2.7 to 4.2 Å (fig. S1). This agreement is especially close in the lowest-energy region of the MFEP (MFEP Location: 20 to 30) and is corroborated by comparable inter-domain distances between the MFEP models and those previously reported (fig. S2).

Despite these similarities, fig. 2 shows that, along the MFEP, the RBD’s Solvent Accessible Surface Area (SASA) scores span the range of values previously assigned to differences between open and closed static structures. The non-biased MD trajectories starting from the apo model span a smaller range of SASA values than those seen in the MFEP, despite an 800-fold longer simulation time. The up RBD’s global conformation is similar to the open spike structures. However, coupling of the global conformations to local changes at key antibody or inhibitor pockets along the MFEP is expected to be significant. This issue is investigated in the next section.

**Figure 2:**
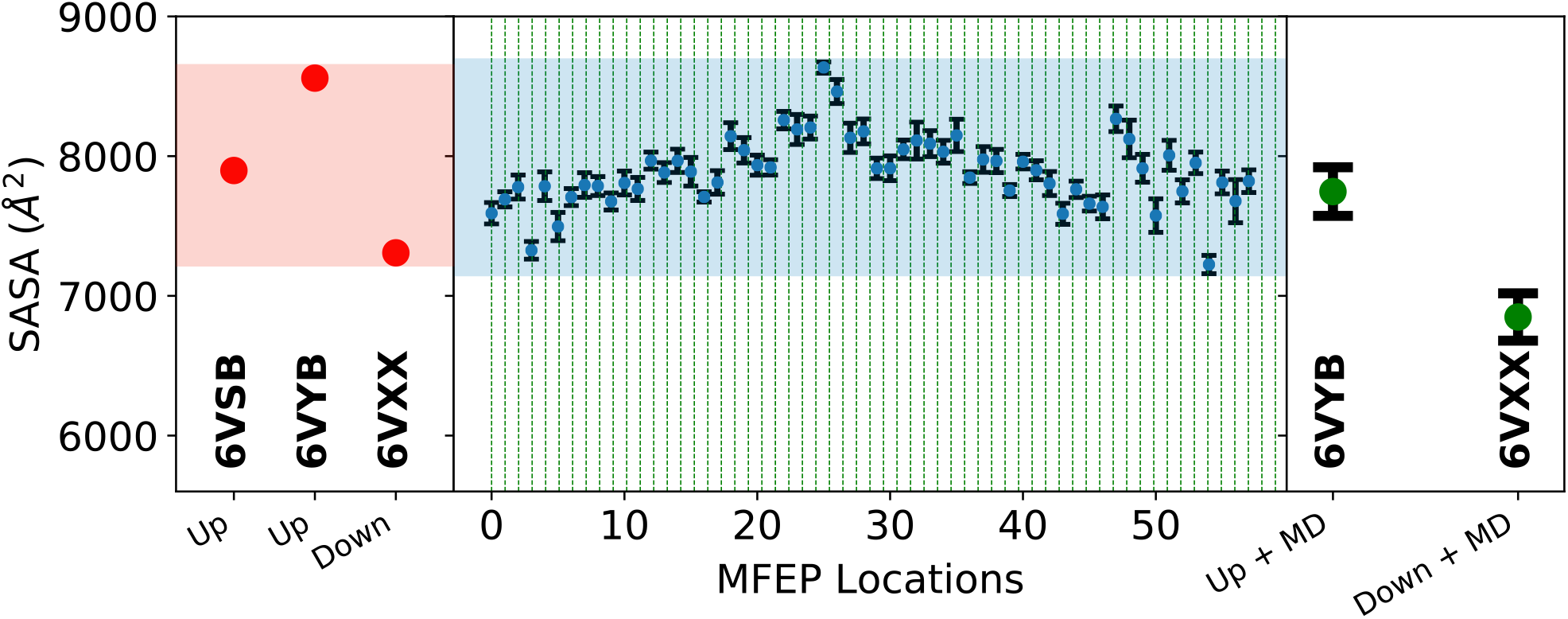
Whole RBD SASA scores calculated for the up RBD (residues 320 to 510). The SASA scores for static structures are represented with red circles in the left panel, the non-biased equilibrium MD trajectories are represented with green circles and error bars in the right panel, and finally each MFEP location SASA scores are represented with blue circles and error bars in the central panel. The error bars show 1 standard deviation from the mean. The red and blue highlighted regions are used to compare the spread of SASA scores of the previously deposited static structures (red shading) with the range of SASA scores seen for all MFEP locations (blue shading).

Globally closed or in RBD-down conformation were not observed in any of the MFEP locations. The distribution of RBD “all-down”, “1-up”, and “2-up” conformations varies across studies (*1,3,32*). Typically, both the “RBD down” and “1-up” conformations have been reported to be in equilibrium, as reflected in the 2:1 to 1:1 population distribution of the 2D images (*1,3*). The lack of RBD-down conformations reflects the absence of equilibrium between the up and the down states in the picked particles used to construct the 6VSB structure. This supports the notion that, in early SARS-CoV-2 spike protein data (*1*), the crown of the spike was over-stabilized to a closed conformation.

### Local conformational motions at binding pockets

The starting structure can have a significant impact on the outcome of computations of ligand protein binding interactions (*33, 34*). Using the MFEP structures derived from experimental data, we are able to investigate the impact of the global conformational changes on individual binding pockets. We choose six previously identified binding pockets: three neutralizing antibodies (*27*); linoleic acid (LA) (*28*); a computationally designed neutralizing mini-protein (*26*); and ACE2 (*4*). By comparing the MFEP structures to apo and holo-structures, we investigate excursions from the reported structures in terms of internal deformations, solvent-accessibility, and the energetics of the individual binding pockets. We also consider the impact of two important glycans N165 and N234 on stabilizing the spike protein complex.

Fig. S4 shows Root-Mean-Square Standard Deviation (RMSD) plots calculated using residues specific to each binding pocket. Despite the highest similarity to 6VSB at MFEP locations 20 to 30 for the global analysis, local variations in RMSD are more pronounced when the binding pockets are aligned by the residues involved in ligand-protein interactions. In fig. 3, we investigate RMSD excursions from the static structure 6VSB aligned using the entire monomer. The pattern observed in the global RMSD analysis recurs, whereby MFEP location 28 has the lowest RMSD to 6VSB. However, each binding pocket deviates from 6VSB. The binding pockets for ACE2 and the mini-protein have high RMSD values at both ends of the horseshoe shaped MFEP, while the binding pockets for LA, AB-EY6A, and AB-CR3022 maintain similar RMSD values in all MFEP locations. The binding pocket for AB-C135 deviates strongly from 6VSB throughout the entire MFEP trajectory. This is discussed further below. In the RMSD space, it is difficult to determine how amenable a pocket is to binding. However, the LA and AB-C135 binding pockets display large differences in RMSD between the apo (6VSB) and holo-structures. The LA binding pocket is distinctly more apo-like, while the converse is true for the AB-C135 binding pocket. To avoid uncertainties in interpreting the local RMSD values, we examine SASA at the local binding-pocket level.

**Figure 3:**
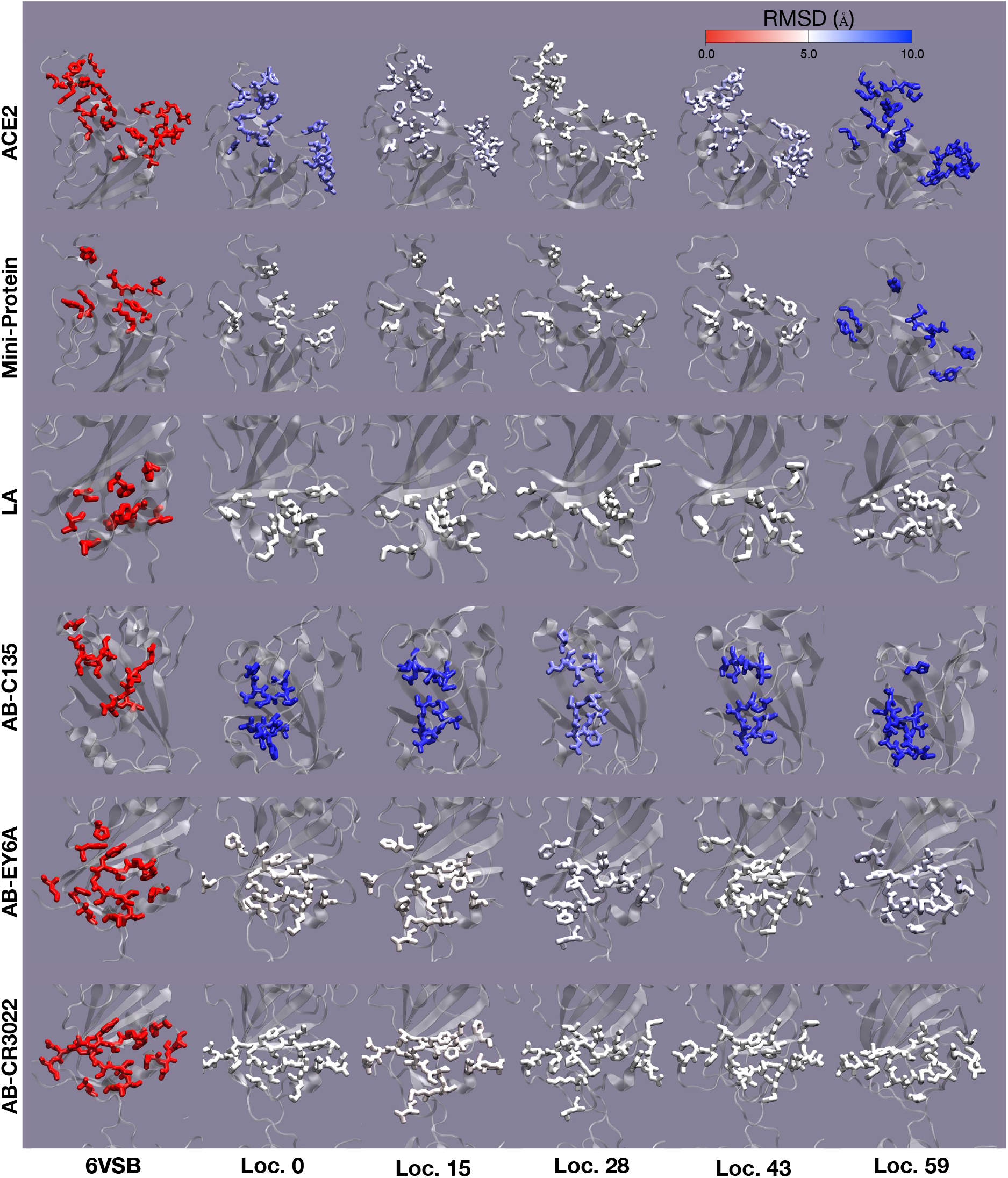
Binding pocket residues colored by RMSD changes across the minimum free energy path. The residues involved in binding are used to calculate the RMSD at MFEP locations 0, 15, 28, 43, and 59 (see Fig. 1) and are shown in a licorice representation. The RMSD values range from 0 to 1 following the color scale at the top of the figure. The structure 6VSB is shown for comparison on the far left.

Fig. 4 shows a clear difference between apo and holo binding pockets. This allows direct comparisons between different MFEP locations, and the tight or loose nature of the binding pockets from holo and apo discrete structures. For five of the six binding pockets, the MFEP reveals a larger or comparable variation in SASA compared to the apo “1-up” RBD structures (6VSB, and 6VYB), the “all-down” RBD structure (6VXX), the holo structure, and 10 microsecond MD ensembles (*29*). For the five binding pockets, where the MFEP locations span the SASA variation or openness of the pocket seen for the static and nonbiased MD structures, the binding mode appears to involve a conformational selection mechanism. For example, the mini-protein binding pocket in fig. 4, at MFEP locations 4, 47, 49, and 54, the calculated SASA scores are commensurate with the holo-structure (PDBID: 7JZU), while the rest of the MFEP locations involve more open structures (higher SASA scores). This suggests that even without the presence of the mini-protein, the binding pocket is able to adopt a tight binding configuration amenable to spike protein neutralization. Interestingly, we observe a key interaction between the LA binding pocket of the “up” RBD and the N234 glycan of the same chain (see fig. 5). At MFEP locations 51 to 59, the N234 glycan on chain A hydrogen bonds with residues 387 (Leu) and 388 (Asn), pulling the binding pocket open, and resulting in the “super-open” states seen in fig. 4 at the same MFEP locations.

**Figure 4:**
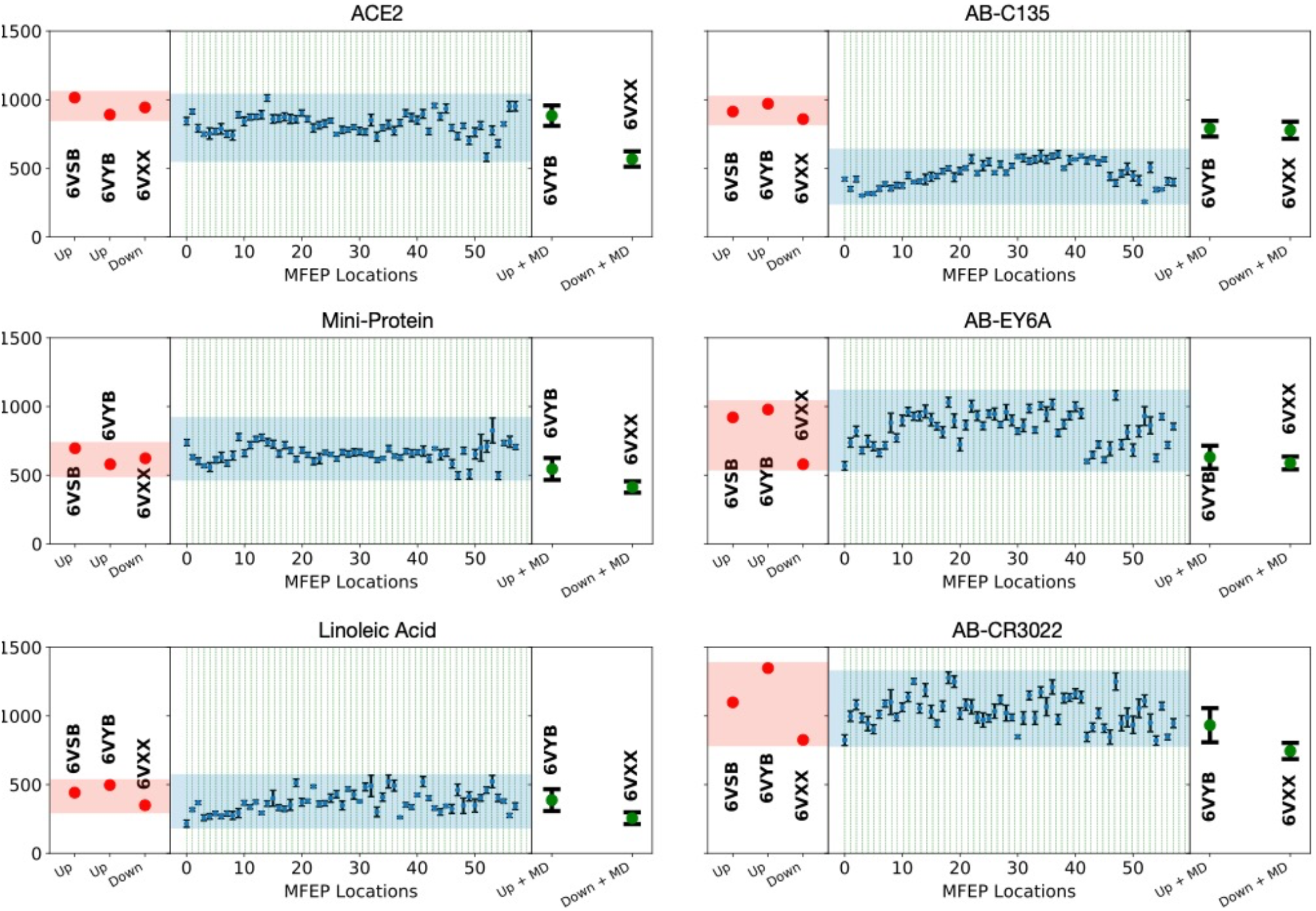
Individual binding pocket SASA scores calculated from only those residues involved in binding. For all six panels, the SASA scores for static structures are represented with red circles, non-biased equilibrium MD trajectories are represented with green circles and error bars, and finally each MFEP location SASA score is represented with blue circles and error bars. The red and blue highlighted regions are used to compare the spread of SASA scores of the previously deposited static structures (red shading) with the range of SASA scores seen for all MFEP locations (blue shading).

**Figure 5:**
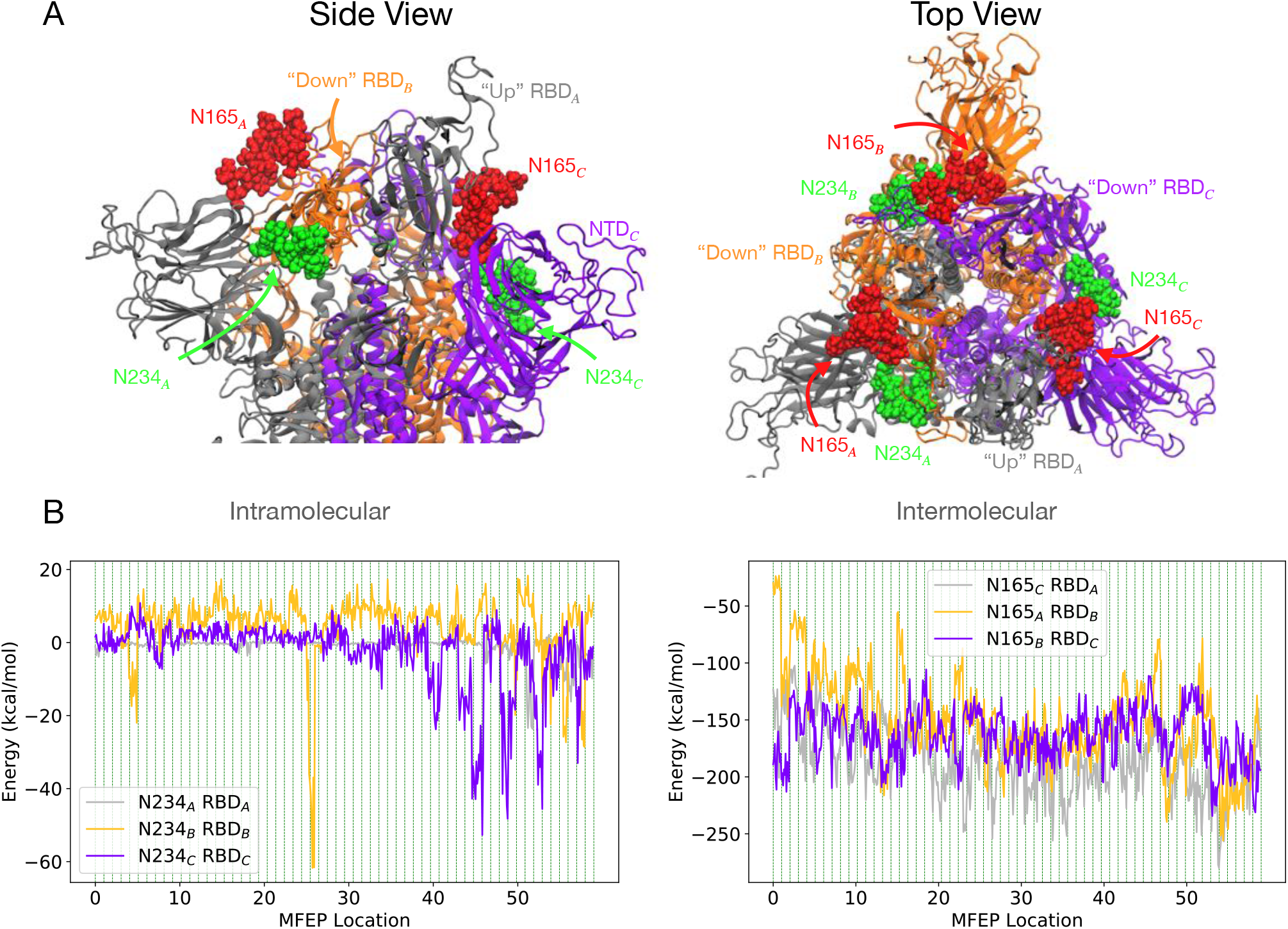
Glycan conformations and interaction energies for all chains. Panel A shows each chain of the spike protein trimer individually colored in a cartoon representation. The glycan chains are shown a red and green surface representation and are labeled accordingly. Panel B shows intramolecular or intermolecular interaction energies of the glycan and the spike protein RBD (residues 330 to 520) for each chain.

The AB-C135 binding pocket has a decidedly more closed conformation (fig. 4). The hydrophobic residues near the AB-C135 binding pocket and at the surface of 6VSB are buried in our simulations, as indicated by the lower SASA score calculated for 6VSB (see fig. S5). This suggests the binding mechanism involves an induced fit. the AB-C135 binding pocket is a “hidden epitope”, indicating that the closed state we find is biologically relevant (*26*). The map resolution surrounding this binding pocket has a marginally lower resolution (see fig. S6), decreasing confidence in the relevance of the observed “super-closed” conformation. The presence of a closed local pocket in an overall up or open spike RBD further suggests that AB-C135 binding entails an induced fit.

So far, we have observed the effect of glycosylation on the LA binding pocket. Fig. 5B shows that N165 has similar pairwise potential energies for all three intermolecular interactions between N165 and the RBD of the counterclockwise chain. The pairwise potential energies for the intramolecular interaction between N234 and the RBD of the same chain show large differences in the ability of the glycan either to stabilize or destabilize the RBD conformation. Both the N234*_B_* RBD*_B_* and N234*_C_* RBD*_C_* pairwise interactions show positive potential energies in many of the MFEP locations due to strong van der Waals interactions. These steric interactions suggest the “down” RBD conformation is destabilized by the presence of N234, as noted by Amaro and colleagues (*26*), albeit only for their intermolecular interactions. The N234*_A_* RBD*_A_* interaction is negligible in most of the MFEP locations, except in the 50 to 59 range of locations. In this region, RBD*_A_* is stabilized by its interactions with N234*_A_*, incidentally creating the “super-open” LA binding pocket discussed above. Finally, fig. S3 shows that comparing SASA values from MD ensembles derived by fitting a single CC with those utilizing the multigrid procedure (i.e., the MFEP locations), the single map CC ensembles do not agree with the MFEP locations at the same CC value. Also, the SASA values for the single CC fitting procedure generally do not span the same range seen for the MFEP locations. This indicates that single-densities based on a single CC do not capture conformations similar to those seen along the MFEP.

## Discussion and Conclusions

An equilibrium sample of biological macromolecules is inherently conformationally heterogeneous, because a range of conformational states can have significant thermally induced occupancies. Conventional cryo-EM data analysis techniques align and average snapshots with similar particle orientations. While this improves contrast in the 2D images, there is a loss of thermodynamic information. Also, static models that fit the averaged 3D reconstruction are ill-suited to describing the system’s conformationally dynamical nature. Starting from such static models, even unbiased MD cannot capture all the low thermally accessible conformational states for a given system due to limited sampling times. By combining data-driven machine learning and MD computations, we have extended the conformational search around the one-up RBD state by making use of experimentally determined energy landscapes. This approach has revealed a broad spectrum of hitherto unobserved iso-energetic conformations associated with RBD binding sites.

The experimentally determined horseshoe shaped MFEP shows it is necessary to move beyond a rigid one-up RBD state picture, because of substantial conformational heterogeneity on local and global scales. The concerted movement of the RBD has regional effects, which entail substantial conformational heterogeneity compared with both apo and holo structures. Nearly all binding pockets analyzed in this study indicate conformational selection mechanism, where specific MFEP locations are more or less favorable to binding. MFEP locations with higher SASA values are interpreted as more amenable to binding, with potential therapeutic implications. The inclusion of a broad spectrum of low-energy conformations can enable new drug discovery routes by considering the extensive conformational flexibility of important binding pockets. By going beyond static structures fitted to heavily averaged maps, our approach reveals the flexible nature of biological macromolecules, which may have implications for novel drugs.

## Methods

### Cryo-EM Data

In this study, we used the cryo-EM images from a previous study (*3*), in which Sars-CoV-2 spike ectodomain residues 1 to 1208 were expressed based on the first reported genome sequence (*35*), adding two stabilizing proline mutations in the C-terminal S2 fusion machinery. Cryo-EM grids were prepared using purified fully glycosylated spike protein. Frozen grids were imaged in a Titan Krios (Thermo Fisher) equipped with a K3 detector (Gatan). Movies were collected using

Leginon at a magnification of x22,500, (*36*) corresponding to a calibrated pixel size of 1.047 A/pixel. A full description of sample preparation and data collection parameters can be found° in (*3*). Motion correction, CTF-estimation, and non-templated particle picking were performed in Warp. (*37*) There were thousands of images in the 631,920 extracted particles which had artifacts e.g. harsh line/boxes. Those images were removed. The remaining 574,324 were imported to CryoSPARC and non-uniform refinement was used to get the orientation of each particle.

### Geometric Machine learning (ManifoldEM)

The details of our data-analytic approach are available at (*6*). In brief, having assigned an orientation to each snapshot, we divide the snapshots to small orientational bins, which we call projection directions (PD). In other words, each PD includes the snapshots which are in closely similar orientations.

We select all the projection directions lying on a great circle around the orientational sphere. Then in each projection direction, we use manifold embedding (*6*) to extract the conformational manifold. We use the two topmost conformational coordinates to describe the conformational manifold If we sort the snapshots along each of these eigenfunctions. We can compile a movie of the conformational changes along those eigenfunctions by using Non-Linear Spectral Analysis (*38*). Fig. 1 shows the first and last frames of the movies (Supplementary Movie 1 and 2) along with the two conformational coordinates. As the movies show, CC1 corresponds to a breathing like motion, while in CC2, the RBD moves from a down position to up.

Integrating the information from all the PDs on a great circle, we compiled 3D conformational movies of electrostatic potential maps along each of these conformational coordinates (Supplementary Movie 1 and 2).

### Map Preparation

We compiled 50 maps (.spi) along each conformational coordinate. The maps along each conformational coordinate were then converted from .spi format to .mtz format with Chimera. (*39*) The .mtz maps were then converted to potentials thresholding the maps at the solvent density peak by utilizing the voltools pot command, which is part of the Voltools Plugin within VMD. (*40*) These potentials were used to guide the spike protein trimer’s dynamics. The maps were thresholded again where the RBD density was weak to increase the magnitude of the potential map gradients for the RBD.

### Molecular Dynamics Simulations

We used models from the CHARMM-GUI Archive - COVID-19 Proteins Library. (*41*) The models were stripped of all water molecules with the molecular visualization program VMD. (*40*) All simulations utilized the generalized Born Implicit Solvent (GBIS) model and Charmm Force Field, (*42*) with the molecular dynamics engine NAMD 2.14b1. (*43*) The simulation parameters are provided in the supplementary NAMD input file.

### Molecular Dynamics Flexible Fitting

MDFF was used to bias simulations and fit atomic models to the extracted conformational coordinates (*24,25*). Noting the medium resolution of our experimental maps, only the backbone of the models was coupled to the density, with the conformations of the remainder of the system (sidechains and glycans) responding to the MD force fields. To ensure that the protein backbone conforms to the density, we constrained the protein by knowledge of its secondary structure, chirality, and cis-peptides; while the less resolved sidechains and glycans continued to refine under the chemical constraints (bonds, angles, dihedrals, and non-bonded interactions) imposed by the CHARMM36m force fields.

### Conformational Coordinate Fitting

To obtain references for the fitting atomic models on the extracted energy landscape, atomic models were fitted to each map for both conformational coordinates. Simulations were biased with two map potentials coupling the singular threshold maps to all backbone atoms except chain A residues 320 to 520, and the doubly thresholded maps were coupled to the backbone atoms chain A and residues 320 to 520. The fitting process occurred in two steps starting from the same starting structure following the cascade-MDFF procedure (*22*). The blurred maps were created with the command voltool smooth in the VMD Voltools Plugin. The first step utilized a 2 *A^°^* Gaussian blurred potential map, while the second step used maps at their original resolution of the maps.

### MFEP Fitting

A movie of molecular motions was compiled along the MFEP of the energy landscape using a so-called “multigrid” fitting procedure (*30, 31*). This procedure enables the construction of atomic models under the influence of two or more density maps. As outlined in Methods, the multigrid procedure is employed within MDFF to enable the fitting of the XY-dimension of the spike protein backbone under the influence of the CC1 maps, while Z-dimensions of these atoms conform to the CC2 maps. A resolution-exchange protocol was concomitantly employed (*22*) to improve the sampling efficiency of the conformational space. Agreement between the single CC and MFEP fitting in terms of radius of gyration is shown in fig. S7.

### SASA Calculations

We used VMD’s measure Plugin to calculate SASA values. The radius sampled around each atom selected was that atom’s radius + 1.4 *Å*. The static models used for comparison were built with VMD using the autopsf Plugin to add H at the physiological pH.

## Acknowledgments

This work was supported by the US National Science Foundation (NSF) under award DBI-2029533. The development of underlying techniques was supported by the US Department of Energy, Office of Science, Basic Energy Sciences under award DE-SC0002164 (underlying dynamical techniques), and by the US NSF under awards STC 1231306 (underlying data analytical techniques) and 1551489 (underlying analytical models). A.S. acknowledges an NSF CAREER award MCB-1942763, and the NIH award R01GM095583. J.V. acknowledges support from the NSF Graduate Research Fellowship Grant 2020298734. We thank J. McLellan and collaborators for providing the cryo-EM dataset, and J. Frank for valuable discussions in the initial stages of the project.

## Author Contributions

AO proposed the study. AO and AS designed the study architecture. All authors contributed to study design. GM extracted the manifold of conformational motions from single particle images, produced the free-energy landscape, and identified the MFEP. JV produced the atomic models for each conformational coordinate and along the MFEP, and analyzed the atomic models. All authors provided critical input and participated in writing the manuscript.

## Data and materials availability

The data supporting the findings of this study are available from the corresponding authors upon reasonable request.

