## Supplemental Material for "Energy Landscape of the SARS-CoV-2 Reveals Extensive Conformational Heterogeneity"

#### Table of Contents

***Supplementary Figures*** ........................................................................................................ ***3***

***Supplementary Movie Captions*** ........................................................................................ ***10***

### Supplementary Figures

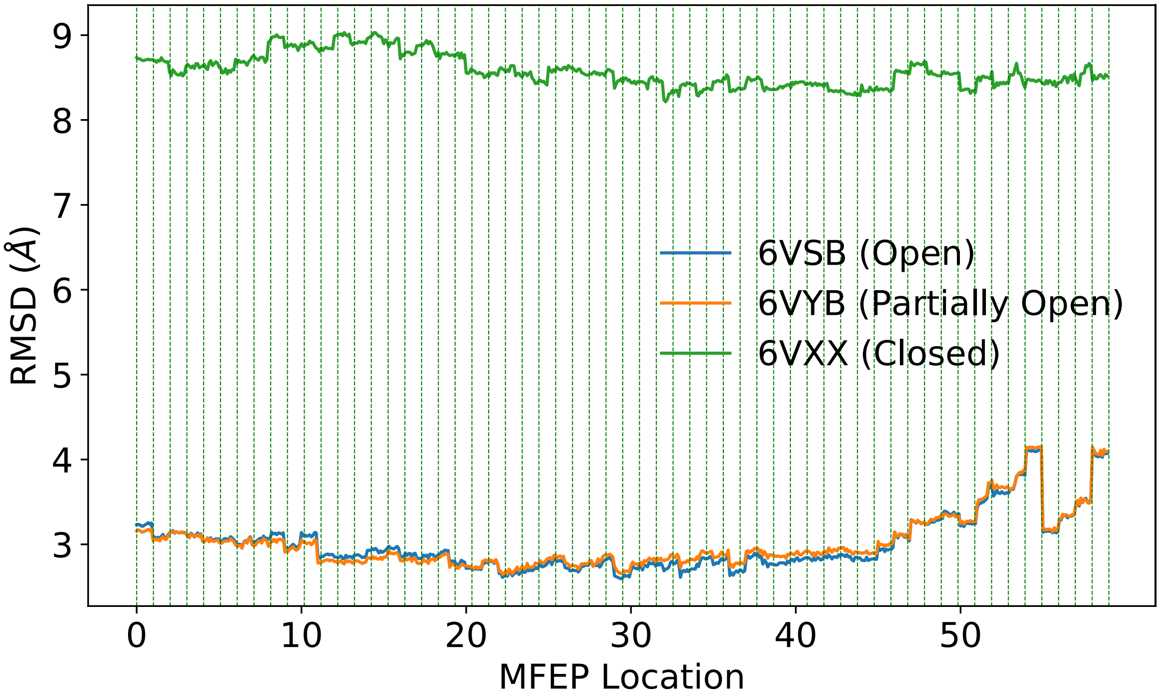


*Figure S1 The RMSD values for ten frames at each MFEP location ensemble. Backbone atoms from chain A were aligned to the reference structure and the RMSD was calculated using the same set of atoms.*

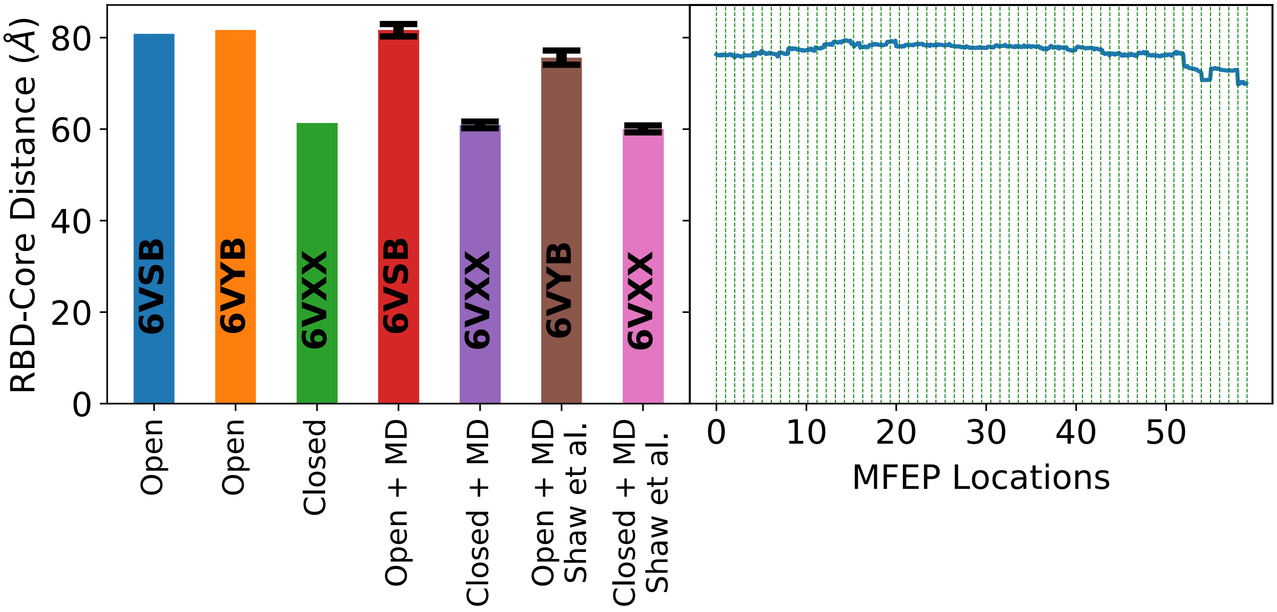


*Figure S2 The distance between the spike protein trimer's center of mass and the center of mass of chain A RBD residues 330 to 510. (left) The distances calculated for the static models (6VSB, 6VYB, and 6VXX). Bars with the axis label "+ MD", signify the mean distance calculated over the trajectory and the error bar represents 1 standard deviation in both directions. The "Open + MD" and "Closed + MD" trajectories are comprised of 100 ns of sampling each in implicit solvent, while the Shaw et al. trajectories are composed of 10 µs of sampling in explicit solvent (20). (right) Shows the distances calculated for ten frames at each MFEP location ensemble.*

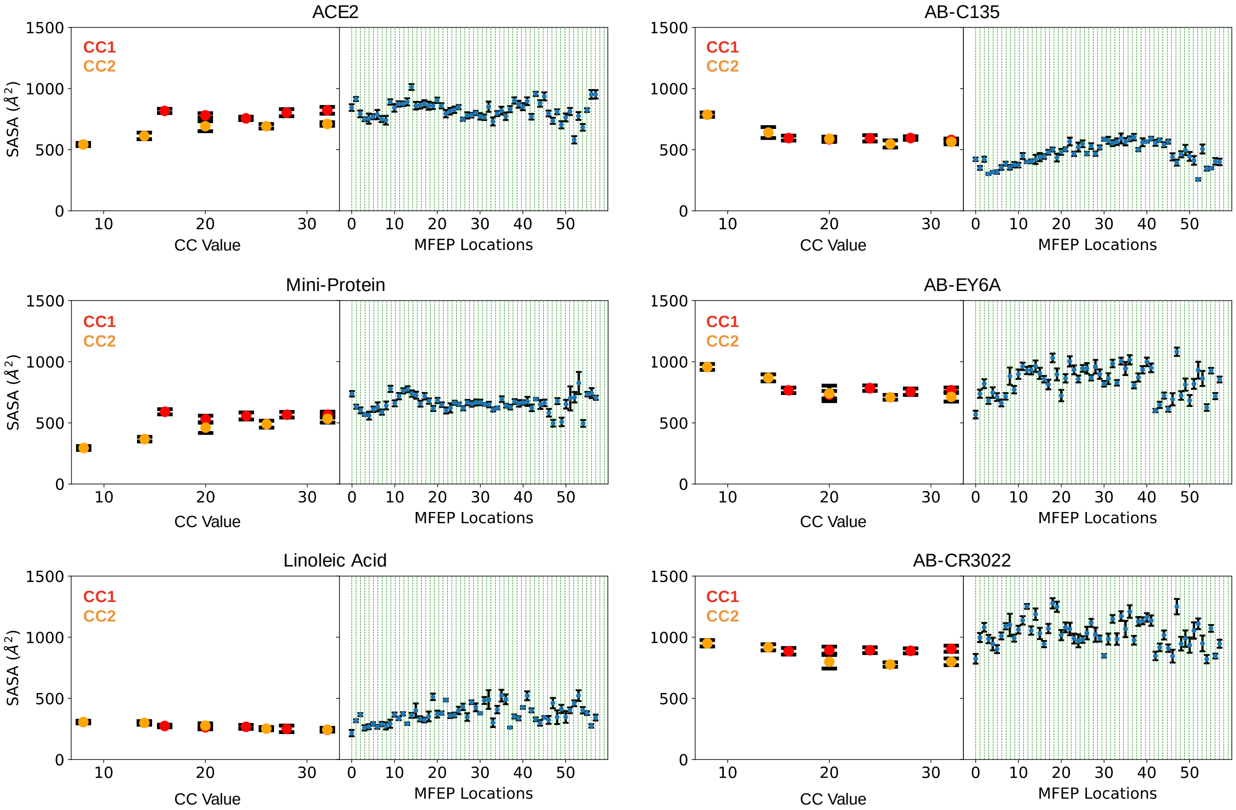


*Figure S3 SASA values of MD ensembles fit to 1 map at a single CC value (left) compared to SASA values at all 59 MFEP locations (right). Circles represent the mean value seen from 10 equally spaced frames coming from the respective ensemble, while the error bars represent 1 standard deviation from the mean in each direction.*

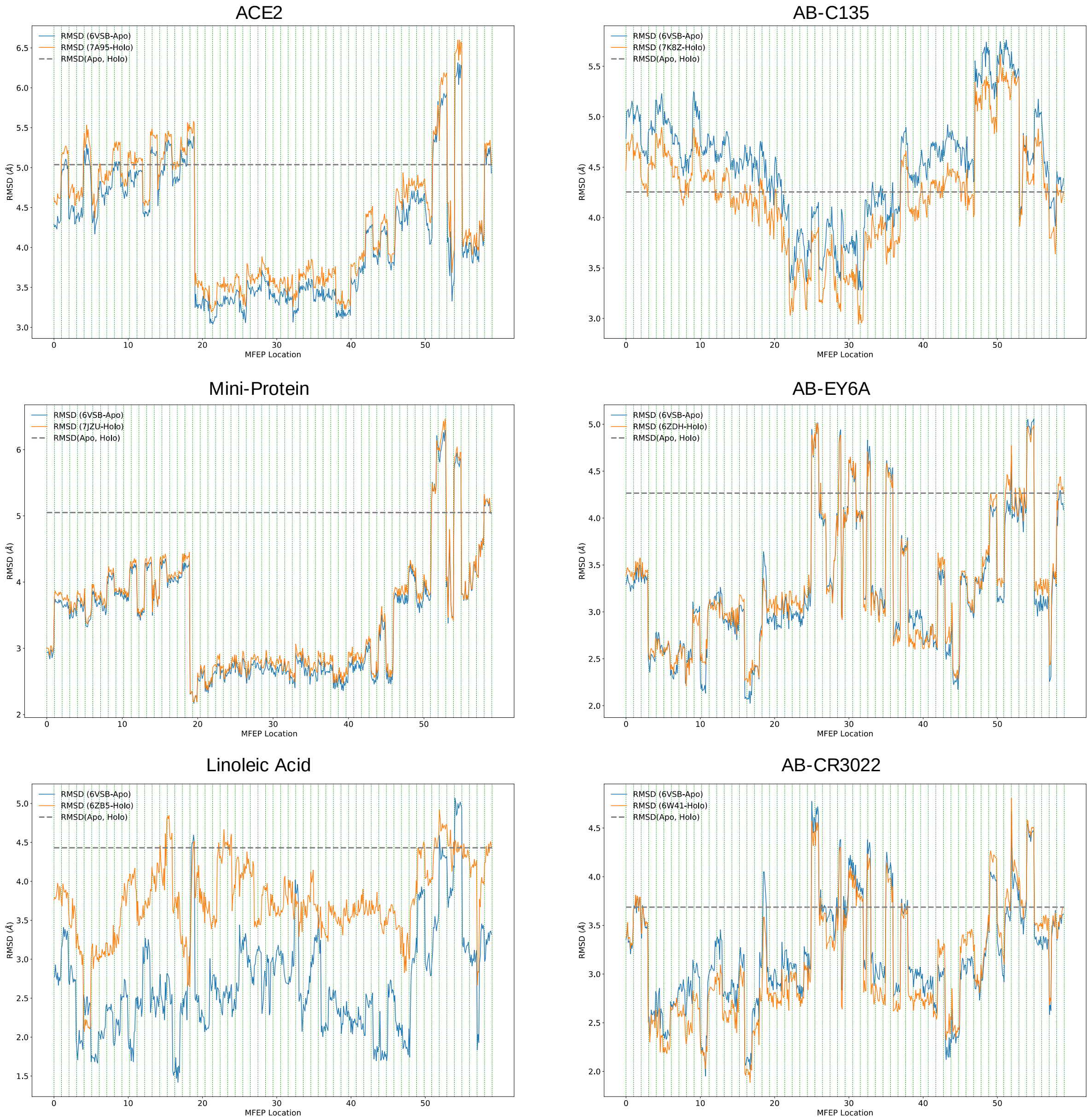


*Figure S4 The binding pocket RMSD between a MFEP location and the static apo (blue line) or holo (orange line) structure. All residues involved in binding were used in the alignment and RMSD calculations.*

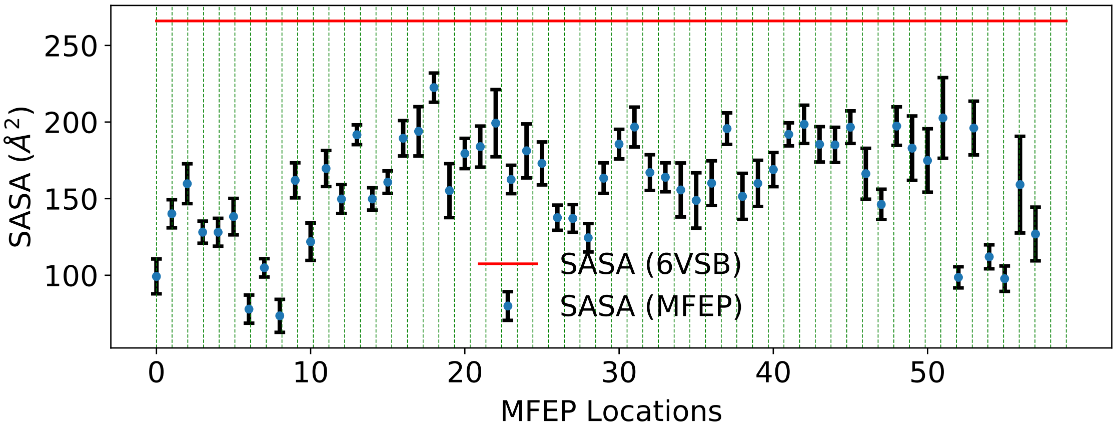


*Figure S5 SASA values of only the hydrophobic residues in the AB-C135 binding pocket for all MFEP locations and 6VSB.*

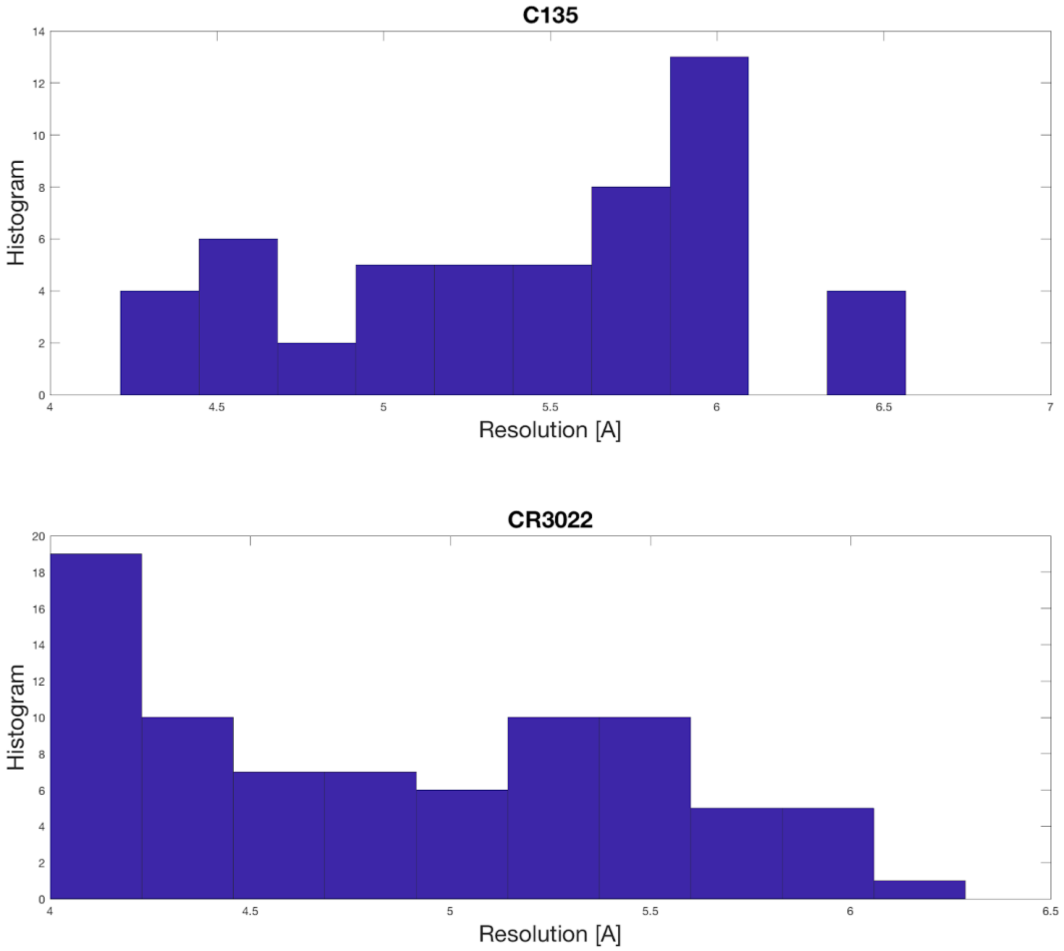


*Figure S6  ResMap (44) results for residues involved in AB-C135 (top) binding and AB-CR3022 (bottom) binding binned by the local resolution around backbone atoms.*

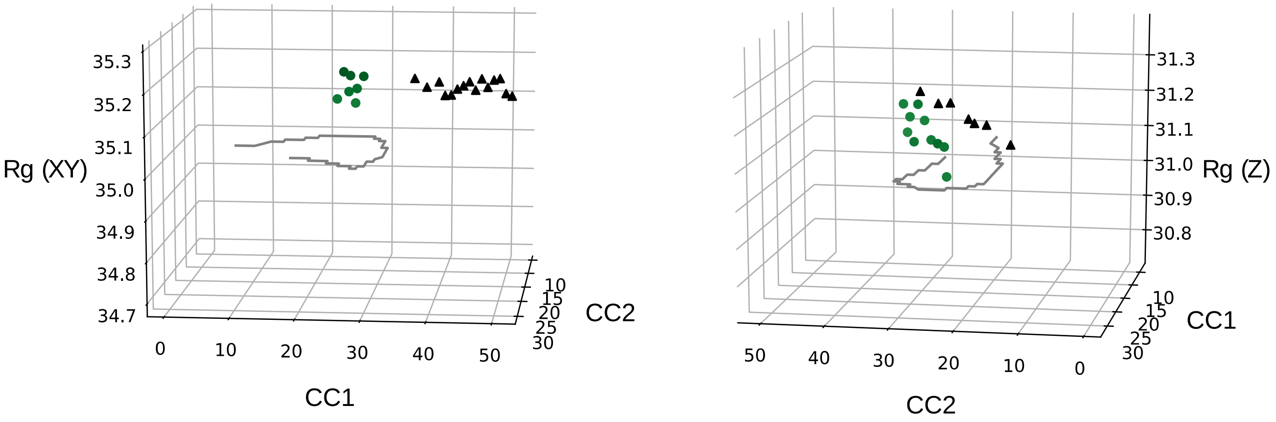


*Figure S7  Comparison of the radius of gyration (Rg) values between single CC fits (black triangles) to those coming from the simultaneous fits of both CCs using the multigrid procedure (green circles). The graph on the left compares the Rg in the XY dimensions of all the MFEP locations and CC1 atomic models. The graph on the right compares the Rg in the Z dimension of all the MFEP locations and CC2 atomic models.*

#### Supplementary Movie Captions

Supplementary Movie 1: CC1 electrostatic potential map movie starting at a CC1 value of 1 and ending at 50.

Supplementary Movie 2: CC2 electrostatic potential map movie starting at a CC1 value of 1 and ending at 50.

Supplementary Movie 3: Energy landscape mapping electrostatic potential maps to atomic coordinates. The current conformation on the right is matched to the movement of the white circle along the MFEP of the energy landscape on the left.
